# Estimating Excitatory and Inhibitory Synaptic Conductances from Spike Trains using a Recursive Bayesian Approach

**DOI:** 10.1101/170878

**Authors:** Mila Lankarany

## Abstract

Inference of excitatory and inhibitory synaptic conductances (SCs) from the spike trains is poorly addressed in the literature due to the complexity of the problem. As recent technological advancements make recording spikes from multiple (neighbor) neurons of a behaving animal (in some rare cases from humans) possible, this paper tackles the problem of estimating SCs solely from the recorded spike trains. Given an ensemble of spikes corresponding to population of neighbor neurons, we aim to infer the average excitatory and inhibitory SCs underlying the shared neural activity. In this paper, we extended our previously established Kalman filtering (KF)–based algorithm to incorporate the voltage-to-spike nonlinearity (mapping from membrane potential to spike rate). Having estimated the instantaneous spike rate using optimal linear filtering (Gaussian kernel), our proposed algorithm uses KF followed by expectation maximization (EM) algorithm in a recursive fashion to infer the average SCs. As the dynamics of SCs and membrane potential is included in our model, the proposed algorithm, unlike other related works, considers different sources of stochasticity, i.e., the variabilities of SCs, membrane potential, and spikes. Moreover, it is worth mentioning that our algorithm is blind to the external stimulus, and it performs only based on observed spikes. We validate the accuracy and practicality of our technique through simulation studies where leaky integrate and fire (LIF) model is used to generate spikes. We show that the estimated SCs can precisely track the original ones. Moreover, we show that the performance of our algorithm can be further improved given enough number of trials (spikes). As a rule of thumb, 50 trials of neurons with the average firing rate of 5 Hz can guarantee the accuracy of our proposed algorithm.

## 1 Introduction

Excitatory and inhibitory synaptic conductances (SCs) provide substantial information about functional mechanism of neuronal sensory responses [1-5]. Most of the past studies in the literature use subthreshold membrane potential to infer SCs [6-12]. Although these methods offer reliable estimates of excitatory and inhibitory SCs at each single trials of recorded membrane potential, they are not capable to infer those SCs, even their average over several trials, from recorded spikes. Thank to recent advancements in neural recordings, scientists can now record spikes from multiple neurons of a behaving animal (in some rare cases from humans) during a specific task. As discussed in different papers in the literature, e.g., see [5] and references therein, having access to underlying SCs will help in better understanding of the mechanism of information processing as well as the corresponding neural circuits. Therefore, inference of SCs only from recorded spikes is the next challenging question in computational neuroscience community.

Among the few papers that address the problem of estimating SCs, that is the average of SCs – not the SCs in single trials, from spikes, Latimer et al [13] proposed a novel extension of popular general linear model (GLM) for spike trains (nonlinear conductance-based model) in order to estimate stimulus-dependent excitatory and inhibitory SCs. They introduced a biophysically inspired point process model that incorporates stimulus-induced changes in synaptic conductance in a dynamical model of neuronal membrane potential [13]. The performance of this method is promising when the stimulus-induced synaptic input is accessible. Moreover, this method is sensitive to precise selection of initial parameters as the log-likelihood function for their model is not concave [13]. It is also to be noted that this method to perform efficient maximum likelihood inference the variability of SCs is neglected; the source of stochasticity is only within the spiking mechanism.

In this paper, we propose a Kalman filtering algorithm that incorporates dynamics of SCs and membrane potential, *states*, as well as instantaneous spiking rate with the same voltage-to-spiking nonlinearity as used in [13], *observation*, to infer excitatory and inhibitory SCs. Hence, our proposed method does not require any information about the external stimulus (note: the identifiability of this problem depends on how precise the subthreshold membrane potential can be estimated from spike trains). As well, the stochasticity of SCs, that is a biologically realistic constraint, is included in our model. Given enough number of spikes based on which an accurate estimate of instantaneous spiking rate is yielded, our method demonstrates robust estimation of excitatory and inhibitory SCs.

## 2 Problem Statement

The instantaneous firing rate of a conductance-based point process can be represented by a nonlinear transformation of its membrane potential [13, 14].

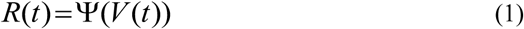

where *Ψ* and *V* are expressed as follows.

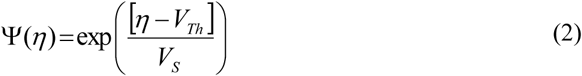

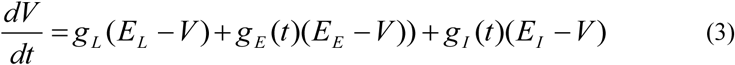

*Ψ* is the voltage-to-spike rate nonlinearity [13] which follows the from proposed by Mensi et al [14]. *V* indicates the sub-threshold membrane potential of a single neuron. In (2), *V_S_* and *V_T_* express the steepness and soft spiking threshold of *Ψ*, respectively. *E_L_*, *E_E_*, and *E_I_*, in (3), are the reversal potentials of the leakage, excitatory and in inhibitory currents, respectively. *gL* is leak conductance. We assume, similar to [8], [9] and [15], that *E_L_*, *E_E_*, *E_I_*, and *g_L_* are known. Following [8, 9, 12], the excitatory and inhibitory SCs, *g_E_* and *g_I_*, can be stated as follows.

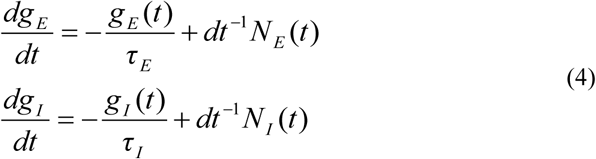

where *τ_E_* and *τ_I_* are excitatory and inhibitory synaptic time constants, and they are known in our simulation studies. And, *N_E_/dt* and *N_I_/dt* (we include *dt^−1^* in (4) to be consistent with the previous definition of synaptic inputs used in [8, 9, 12]) indicate excitatory and inhibitory synaptic inputs, respectively.

Here, we emphasize that the instantaneous firing rate, *R(t)*, is obtained, in this paper, from recorded spikes where an optimal (Gaussian) kernel [16] linearly maps spikes to *R(t).* Therefore, the higher number of spikes the more accurate *R(t)*. As the firing rate of a single neuron is not sufficiently large, we assume that spike trains from multiple trials (for the same neuron) are available. Hence, we state our problem to estimate the average (shared) of SCs of a single neuron expressed by (3) from ensemble of spikes during multiple (repeated) stimulation trials. The external stimulus (it is different from synaptic inputs) is not available for calculating SCs, hence our algorithm is developed to infer SCs from spikes only. This problem is not identifiable in a general perspective. However, assuming that the neurons are homogeneous and their biophysical parameters are known, one can interpret that the spike rate is reproducible if and only if a certain external stimulus is presented. Figure 1 illustrates the schematic of our problem.

**Figure 1:**
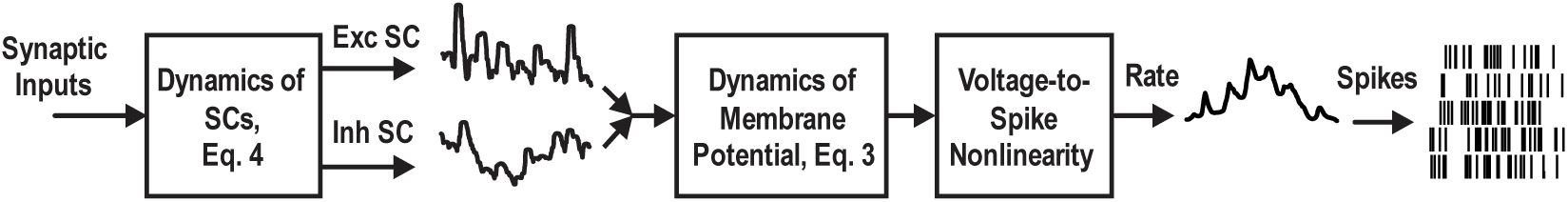
Schematic of spike generation based on excitatory and inhibitory SCs.

By incorporating Eq (1-4), we express the dynamical system underlying the instantaneous firing rate of an ensemble of neurons (Figure 1) as a state-space model.

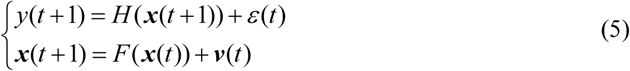

where ***x***(*t*) and *y*(*t*) indicate the state vector (including the subthreshold membrane potential and synaptic conductances) and the observation (instantaneous firing rate) at time *t*, respectively. Functions *F* and *H* are the transition and observation functions, respectively, and ***v***(*t*) and *ε*(*t*) are the system noise (comprising synaptic inputs) and the observation noise, respectively. An explicit discrete form (5) is presented as follows (see [8] for details).

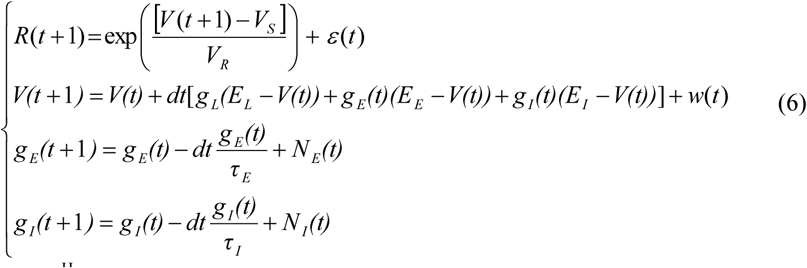

where [*V*(*t*), *g_E_*(*t*), *g_I_*(*t*)]^H^ denotes the state vector ***x***(*t*), in (5), and *R*(*t*) is the observation at time *t*, i.e., *y*(*t*) in (5). The transition function *F*, is given by

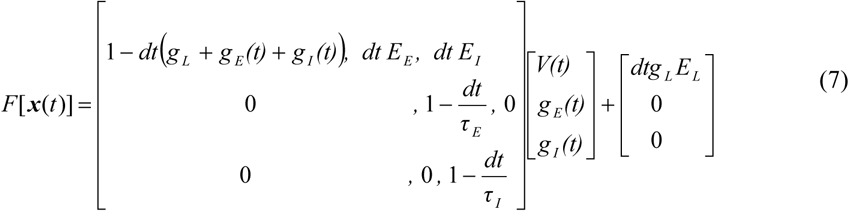

And, the distribution of the system noise (dynamical noise) ***v*** (*t*)=[*w*(*t*), *N_E_*(*t*), *N_I_*(*t*)]^H^ is given by

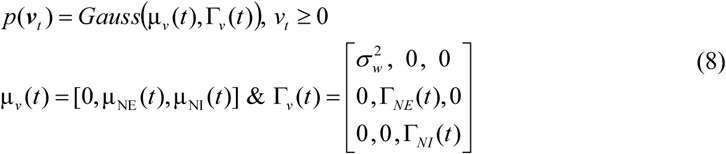

where *N_E_* and *N_I_* describe excitatory and inhibitory synaptic inputs, respectively. Finally, the observation function *H* is represented by:

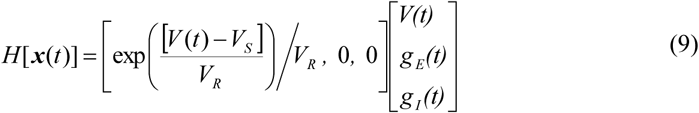

## 3 Proposed Algorithm

In this section, we derive an extended Kalman filtering (EKF) followed by expectation maximization (EM) algorithm for (6) to infer the average excitatory – inhibitory SCs (*g_E_*(*t*) and *g_I_*(*t*) in (6)) of a single neuron in response to a repeated stimulus.

### 3.1 Kalman Forward/Backward Filtering

As the transition and measurement functions, in (6) or its equivalent in (5), are nonlinear, we employ EKF that uses the first-order Taylor linearization of the nonlinear process and measurement model to derive the underlying prediction–correction mechanism. We calculate the state estimate E{***x***(*t*)[*y*(0:*t*)} and state correlation matrix E{***x***(*t*)***x***(*t*)^H^ | *y*(0:*t*)} in the forward filtering step and E{***x***(*t*)|*y*(0:*T*)} and E{***x***(*t*)***x***(*t*)^H^ | *y*(0:*T*)} in the backward filtering (smoothing) step using the KF approach for the observed instantaneous spike rate. Here, E{⋅} stands for the expected value. More details of KF forward and backward filtering can be found in [8].

### 3.2 Inferring Statistical Parameters via Expectation Maximization

EM algorithm is derived for (6) to infer the statistics underlying excitatory and inhibitory synaptic inputs from the instantaneous firing rate which is already calculated from all recorded spikes. Specifically, EM algorithm (see Eq. 8) infers the time-varying mean (***μ**_v_* (*t*)) and the variance of the states (*σ*^2^_*w*_, *Γ_v_*(*t*)), and the variance of the observation noise (*σ*^2^_*ε*_). Again, it is to be noted that these statistics reflect the average (shared) excitatory and inhibitory synaptic input a single neuron receives in response to repeated trials of an identical stimulus. Incorporating all estimated statistics of the state estimates (mean and correlation matrices) in Kalman filtering steps, the EM algorithm can be easily derived by maximizing the logarithm of the joint probability of the states and observation (*X*and *Y* denote the entire samples of ***x*** and*y* over time, respectively) as follows:

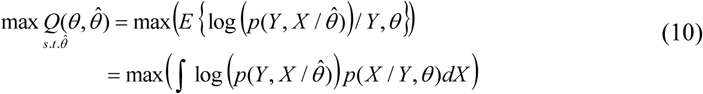

To solve (10), we can write:

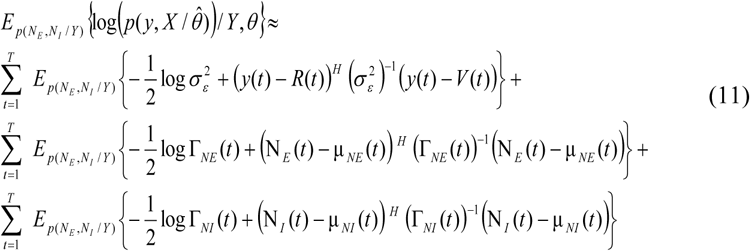

where we assumed that the variance of membrane potential is negligible. By taking derivative of (11) with respect to the mean and variance of the synaptic inputs (statistical parameters are assigned to [*μ_E_*, *μ_E_*, Γ_*E*_, Γ_*I*_] where the variance of membrane potential observation noise are not included), we can update these statistical parameters (see Appendices of [8] for full derivations). Inferring all parameters, we can initialize the next iteration of the recursive algorithm (see Figure 1 of [12] for better understanding of the recursive algorithm). The algorithm continues until there is no significant change (<5% increase in likelihood function) between two consecutive iterations. The statistical parameters of the excitatory and inhibitory synaptic inputs are estimated using B-spline method (see [8, 9] for details) as follows:

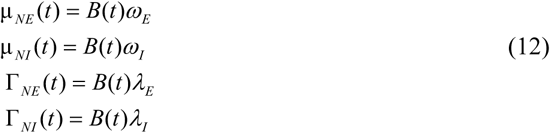

where,

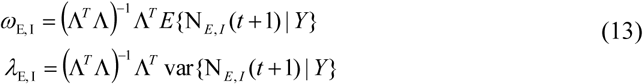

Matrix Λ consists of 50 spline basis functions. Here, *μ_NE_*, *μ_NI_*, *Γ_NE_*, and *Γ_NI_* are the mean (μ) and the variance (Γ) of the excitatory and inhibitory synaptic inputs. To calculate the conditional mean and variance of synaptic inputs, we have:

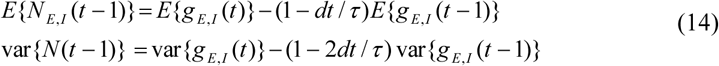

where (14) is derived from (6), i.e., *g*_*E*,*I*_ (*t* +1) = (1 – *dt* / *τ*_*E*,*J*_)*g*_*E*,*I*_ (*t*) + *N*_*E*,*I*_ (*t*). Given the observation over whole recoding time [*0:T*], the conditional mean and variance of synaptic inputs can be then calculated from those of SCs (already available from KF step). We have:

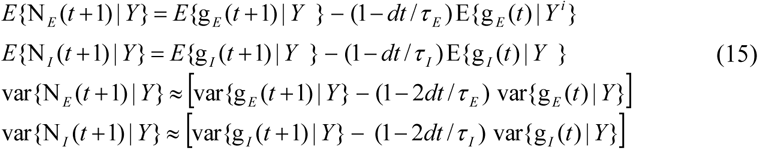

As pointed out earlier, our algorithms operate in a recursive manner using the inferred statistical parameters (of synaptic inputs, see Figure 1) to update the estimates of SCs.

## 4 Simulation Results

Two scenarios are considered, in this paper, to test the accuracy of our proposed algorithm in estimating the average excitatory and inhibitory SCs from recorded spike trains. *Scenario I* is, in fact, a proof of principle where the instantaneous spike rate is directly calculated from subthreshold membrane potential using voltage-to-spike nonlinearity (Figure 2). In *Scenario II*, we use leaky integrate and fire (LIF) model to generate spikes (Figure 3). Hence, first we demonstrate the practicality of our algorithm when there is no uncertainty in the observation, i.e., the source of variabilities is in SCs. Second, we analyze the performance of our proposed algorithm when observation (spike rate) is obtained from spikes generated through multiple trials.

**Figure 2:**
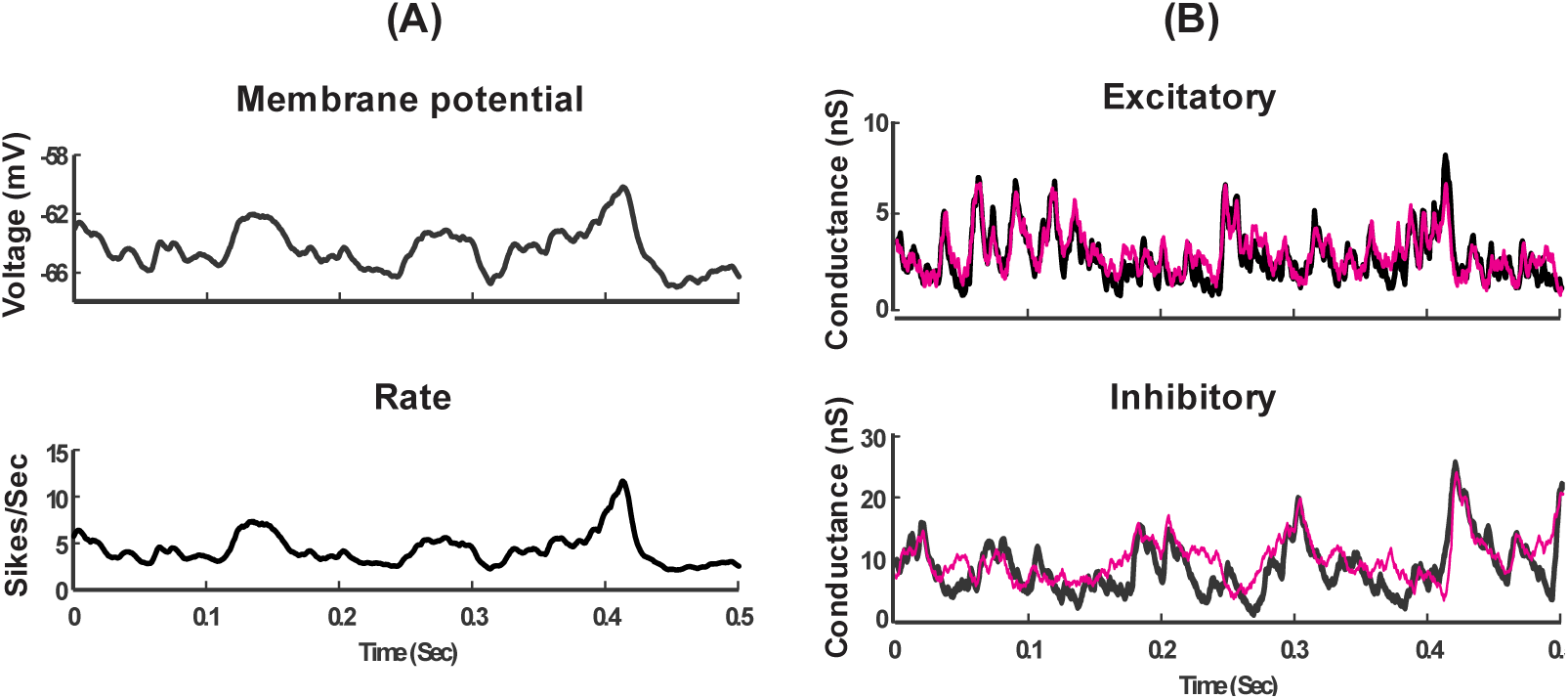
Estimating excitatory and inhibitory SCs in Scenario I (rate function (observation) is a nonlinear transform of subthreshold membrane potential, VT = −70, Vs = 4). (A) Average subthreshold membrane potential (top) and rate function (bottom). (B) Estimated (pink) vs. original average (black) of excitatory (top) and inhibitory (bottom) synaptic conductances.

**Figure 3:**
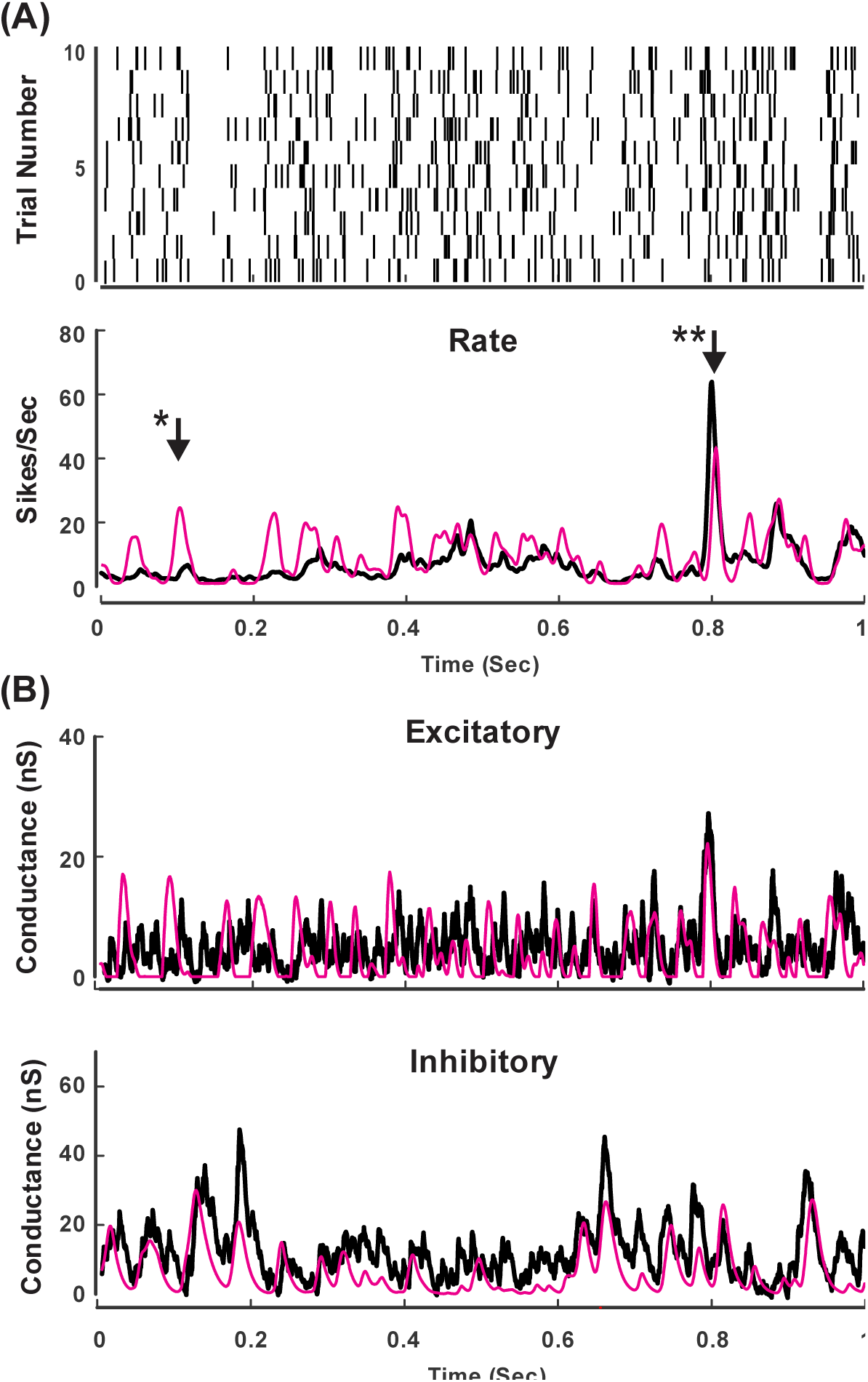
Estimating excitatory and inhibitory SCs from spike trains generated by LIF neuron model (*N* = 50, average firing rate = 4.89 Hz). Note: each trial index (panel A (top)) includes superimposed spikes from five trials, hence all spikes from trials, *N* = 50, is shown in this raster plot. (A) Raster plot of spikes (top), and estimated vs. original rate function (bottom). Here the original rate function is obtained by the nonlinear transformation of subthreshold membrane potential (VT = −70, Vs = 4), and the estimated one is obtained by linear filtering the spikes with an optimal Gaussian kernel [16]. (B) Estimated (pink) vs. original average (black) of excitatory (top) and inhibitory (bottom) SCs. The arrows in panel (A) demonstrate two timing points where the quality of estimated SCs is different (further discussed in the main body).

The parameters underlying the subthreshold membrane potential is listed as follows. The leak potential is E_L_ = –60 mV and the membrane time constant is 1/g_L_ = 20 msec (similar to [8], all conductances are normalized to the cell capacitance). Excitatory and inhibitory reversal potentials are E_E_ = 0 mV and E_I_ = –80 mV, respectively. Excitatory and inhibitory synaptic time constants are **τ**_E_ = 3 ms and **τ**_I_ = 10 ms, respectively. Sampling time is 2 msec and the total simulation time is 2 second. And, a white Gaussian observation noise of standard deviation (*std*) 0.1 mV (which is almost negligible when compared to *std* used in [8, 12]) is added to the membrane potential at each time step.

Excitatory and inhibitory synaptic inputs (*N_E_* & *N_I_*) in each trial are randomly generated by a Poisson distribution given the common mean for each excitatory and inhibitory currents (*μ_NE_* & *μ_NI_*). These time varying statistics are modeled by Ornstein–Uhlenbeck (OU) processes (the absolute white noise (non-negative) is filtered based on excitatory and inhibitory time constants). Then, SCs and the resulting membrane potential traces are generated according to (6). In *Scenario I*, the instantaneous spiking rate is also obtained by (6), and used as observation. In *Scenario II*, this rate is calculated by filtering the spikes by an optimal Gaussian kernel whose width is estimated by the method proposed in [16]. The characteristic of voltage-to-spike nonlinearity is the same as that used in [13] where *V_Th_* = −*70* mV and *V_S_* = *4* mV.

In LIF model, spike time *t_sp_* is recorded when *V(t)* in (3) crosses the spike threshold (-20 mV here) from below. After each spike we included a dead-time [17] of 1 msec before the voltage restarts from resting potential. It is to be noted that we add, in each trial, an independent additive noise to the neuron (OU process: mean = 400 pA, std = 100 pA, time constant = 5 msec) to keep the firing rate between 3-7 Hz.

Figure 2 shows the results of *Scenario I*. Spike rate and membrane potential are shown in Figure 2.A. Given the spike rate, we apply our proposed algorithm to infer the excitatory and inhibitory SCs. As we can see in Figure 2.B, both excitatory and inhibitory SCs are accurately estimated. Note: excitatory SCs are estimated better because the driving force inhibitory SC is small (see [9, 13] for further discussions).

Figures 3 shows the performances of our proposed algorithm in estimating the average excitatory and inhibitory SCs from recorded spike trains (*N_trial_* = 50). In this figure, panel (A) shows the raster plots and corresponding spike rate (true: black, observation (estimated by Gaussian kernel): pink). As can be observed, the estimated spiking rate is relatively fitted to that obtained by nonlinear transformation of subthreshold membrane potential (black). Panel (B) demonstrates the estimated SCs (pink). Both excitatory SCs are estimated with good precision, i.e., the inferred one can almost track all different amplitudes of the original one (although inhibitory SC is slightly underestimated due lower driving force). As clear in Figure 3, the performance of the estimated SCs is constrained by the accuracy of the estimated spike rate. The better the latter is the better the former will be resulted. For example, two different time points are shown in the spike rate in Figure 3. Panel (A) by arrows. The arrow followed by a *single* ^∗^ indicates a case where the estimated SCs do not completely track the original ones. This is because the underlying spike rate is not well fitted to the original rate, i.e., the nonlinear mapping of the subthreshold membrane potential. Nevertheless, the arrow followed by a *double* ^∗^ indicates a case where the estimated SCs accurately track the original SCs (though the inhibitory SC is still underestimated, see above-mentioned reason). It is obvious that such reliable estimates of SCs occurred as the result of accurate estimation of the spike rate. Furthermore, we observed that the performance of estimated SCs enhances for larger number of trials.

To quantify the accuracy of our algorithm for different number of trials, we quantify the total normalized error of excitatory and inhibitory SCs as follows.

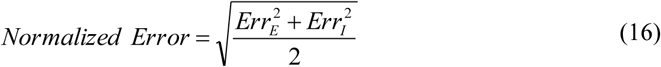

where 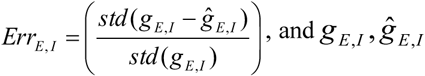indicate the true and estimated SCs, respectively.

Err E,I is a normalized error measure for estimated excitatory and inhibitory SCs. Figure 4 demonstrates how the performance of our proposed algorithm varies with number of recorded trials. In this figure, the average of normalized error is calculated over five different simulations, each of which comprise various number of trials as shown in the figure (the total average firing rate for all neurons is 4.89 Hz). The small arrow in **Figure 4** indicates trial number = 50 where the normalized error is small enough to rely on the estimates. In fact, we note, as a rule of thumb, that 50 trials of spikes (average firing rate of 5 Hz) is a minimum number of trials to reliably infer excitatory and inhibitory SCs.

**Figure 4:**
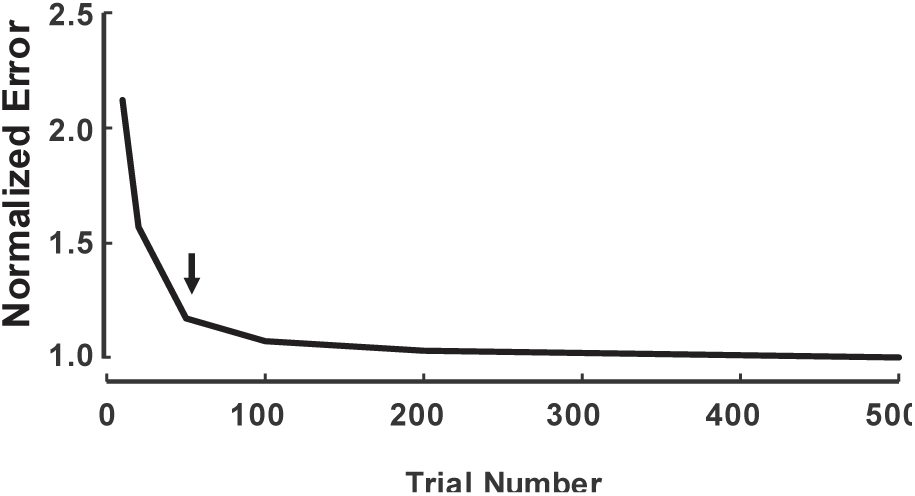
The performance of our proposed algorithm for different number of trials. The normalized error (see (16)) is calculated for different number of trials. The results are the average over five different simulations. The small arrow indicates the minimum number of trials required for a proper performance of our algorithm.

## 5 Summary

We proposed a recursive KF-based algorithm to infer the excitatory and inhibitory SCs from recorded spikes. The instantaneous spike rate was considered as the observation where the voltage-to-spike nonlinearity maps the dynamics of subthreshold membrane potential to that rate. Unlike other related works such as [13], our method does not require to have access to the external stimulus. As well, different sources of stochasticity (unlike [13] that only considers the spike variability) were included in our method. The accuracy of the proposed algorithm was validated through various simulations. Using spikes generated by LIF neuron model, we found that our algorithm works properly given at least 50 trials of a 5Hz neuron (as a standard firing rate for real neurons). Furthermore, we found that the performance of the estimated SCs improves for larger number of trials as the estimated spike rate can better track the actual rate. Consistent to [13, 14], the characteristics of voltage-to-spike nonlinearity are fixed in our model, i.e., *VT* = −70, *Vs* = 4. However, these parameters can be optimized using a likelihood function that directly maps the neuron dynamics to the spike rate. Further investigation in this direction builds the topic of our future studies. Finally, we believe that this work opened a new insight for designing algorithms that can infer both excitatory and inhibitory SCs solely from recorded spikes.

